# Surfactant-Assisted Colorimetric Signal Enhancement in Paper-Based Glucose Sensing

**DOI:** 10.64898/2026.09.21.753053

**Authors:** Dain Kang, Jongho Lee, Minwoo Bae, Seokbeom Roh, Da Yeon Cheong, Hyungbeen Lee, Taeha Lee, Gyudo Lee

**Affiliations:** Department of Biotechnology and Bioinformatics, Korea University, Sejong 30019, South Korea; Interdisciplinary Graduate Program for Artificial Intelligence Smart Convergence Technology, Korea University, Sejong 30019, South Korea; Digital Healthcare Center, Sejong Institute for Business and Technology, Korea University, Sejong 30019, South Korea; Department of Digital Healthcare Engineering, Korea University, Sejong 30019, South Korea

**Author notes:** Corresponding authors: T.L.; G.L. These authors equally contributed to this work (Dain Kang, Jongho Lee).

**Keywords:** paper-based analytical device, surfactant, Tween 20, colorimetric glucose sensor

## Abstract

Paper-based colorimetric sensors offer a low-cost and accessible platform for point-of-care (POC) analysis, but enzyme activity loss during coating and drying can weaken analytical signals and require high enzyme loadings or complex immobilization procedures. Although surfactants are widely used to improve wettability in paper-based assays, their potential contribution to colorimetric performance beyond these effects remains unclear. Here, we investigated surfactant-assisted colorimetric signal enhancement in a glucose assay implemented on a 96-puddle paper plate (96-PPP) and identified Tween 20 as the most effective surfactant. Its effect on detection performance became more pronounced as glucose oxidase (GOx) loading decreased; at 0.1 mg/mL GOx, Tween 20 lowered the limit of detection (LoD) from 0.113 to 0.034 mg/mL (approximately 3.3-fold) over a working range of 0–5 mg/mL, despite no statistically significant change in the measured contact angle at this loading. Tween 20 had no appreciable effect on the reaction in solution but preserved 95% of the apparent reaction rate constant after drying, compared with 11% without it, and atomic force microscopy (AFM) revealed a more dispersed dried enzyme morphology on mica. Tween 20-containing sensors also showed slower signal decay during repeated wetting–drying cycles and thermal stress, retained 77% (vs 26%) of the response at 400 mM NaCl, and exhibited within-PPP and between-batch coefficients of variation (CVs) below 10% (vs 12.3–19.5%), while maintaining glucose selectivity over potentially interfering molecules. These results indicate that Tween 20 enhances paper-based glucose sensing beyond wettability, in part by retaining enzyme cascade activity during drying, although the contributions of the individual enzymes and the underlying mechanism remain to be established.

## 1. Introduction

Paper-based analytical devices (PADs) have attracted considerable attention as point-of-care (POC) diagnostic platforms because of their low cost, portability, ease of fabrication, and low sample and reagent requirements.^1–3^ Cellulose-based substrates may also reduce reliance on plastics and other nonbiodegradable materials used in disposable diagnostic devices. Beyond these practical advantages, advances in paper electronics and lab-on-paper technologies have highlighted the functional versatility of cellulose-based materials in device fabrication and biosensing.^4–6^ These developments illustrate the potential of paper-based materials to support diverse analytical functions for POC applications. Among the biomarkers targeted by these platforms, glucose is a representative biomarker for the diagnosis and management of diabetes, for which frequent monitoring is essential.^7–10^ Accordingly, glucose detection remains one of the most extensively investigated applications of paper-based POC sensing platforms.^11, 12^

Paper-based glucose sensors employ various detection modalities, including electrochemical, fluorescence-based, and colorimetric methods, coupled with enzymatic or nonenzymatic catalytic reactions.^13–15^ Among these approaches, colorimetric detection is particularly well suited to POC testing because the results can be interpreted visually or quantified by simple image analysis without sophisticated instrumentation.^16, 17^ Although nonenzymatic catalytic systems can offer high physicochemical stability,^18, 19^ enzyme-based colorimetric sensors provide high substrate specificity through intrinsic molecular recognition and enable straightforward assay configurations under mild aqueous conditions using commercially available enzymes with standardized activity units.^20–22^

Nevertheless, enzymes are sensitive to their physicochemical environment, and their activity may decrease during deposition and drying on paper substrates owing to processes such as interfacial adsorption, conformational rearrangement, and intermolecular aggregation.^23–25^ Importantly, many paper-based glucose assays employ a glucose oxidase–horseradish peroxidase (GOx–HRP) enzyme cascade rather than a single catalytic reaction. Because the analytical response depends on the sequential function of both enzymes, partial inactivation of either catalytic step during drying can reduce cascade efficiency and ultimately decrease the colorimetric signal. Increasing the enzyme loading can compensate for this activity loss but increases reagent costs, whereas immobilization methods such as covalent conjugation and cross-linking introduce additional fabrication steps and may alter enzyme activity.^26, 27^ Therefore, a simple strategy is needed to preserve the functional integrity of enzyme cascade reactions during drying while retaining the low cost and ease of fabrication of paper-based enzymatic sensors.

Surfactants offer a straightforward means of addressing this challenge. In paper-based assays, surfactants have primarily been investigated for their ability to reduce surface tension and improve substrate wettability, thereby facilitating sample spreading, reconstitution of dried reagents, and mass transport of reactants.^28, 29^ Nonionic surfactants can preferentially occupy air–liquid and solid–liquid interfaces, thereby reducing interfacial protein adsorption. They may also interact with exposed hydrophobic regions of proteins and limit aggregation or inactivation.^30–32^

In enzyme cascade systems, preserving the activity of each catalytic component is particularly important because partial deactivation at any step can limit the overall reaction efficiency and colorimetric output. Thus, the performance enhancement observed in surfactant-modified paper-based enzymatic sensors may arise not only from improved wettability but also from the preservation of enzyme activity during drying and subsequent rehydration. Although the role of surfactants in protein stabilization is well established in solution and in formulation science, their use in paper-based assays has primarily focused on regulating fluid transport and sample spreading. Systematic comparisons of different surfactants in dried paper-based enzymatic assays, particularly those that distinguish enzyme-stabilizing effects from wettability-driven signal enhancement, remain limited.

Herein, we report a surfactant-assisted strategy for enhancing the colorimetric response and detection performance of a GOx–HRP-based glucose assay implemented on our previously developed 96-puddle paper plate (96-PPP) platform.^33, 34^ Comparative screening and concentration optimization identified Tween 20 as an effective additive, particularly when enzyme loading was reduced. Atomic force microscopy (AFM) revealed changes in the distribution of the dried enzyme mixture, and a dry–rehydrate experiment showed that Tween 20 preserved the apparent reaction rate of the GOx–HRP cascade after drying. Stress testing further demonstrated improved retention of the colorimetric response under wetting–drying, thermal, and high-ionic-strength conditions. Collectively, these findings indicate that surfactant addition provides a simple means of improving the performance of dried paper-based enzymatic sensors without complex immobilization procedures.

## 2. Results and Discussion

### 2.1 Principle of the Surfactant-Assisted Colorimetric Glucose Assay

Glucose detection was implemented through a cascade reaction in which GOx oxidizes glucose to generate hydrogen peroxide (H₂O₂), which is subsequently used by HRP to oxidize the colorimetric indicator 2,2′-azino-bis(3-ethylbenzothiazoline-6-sulfonic acid) (ABTS) to its blue-green radical cation (ABTS•⁺) (Figure 1a).^20^ Because coating and drying can compromise enzymatic performance on paper, we hypothesized that surfactant addition could improve wetting behavior while mitigating the loss of enzymatic performance associated with drying (Figure 1b). Before the main experiments, the coating sequence, deposition method, and buffer composition were optimized to establish the baseline assay conditions (Figure S1 and S2).

**Figure 1.**
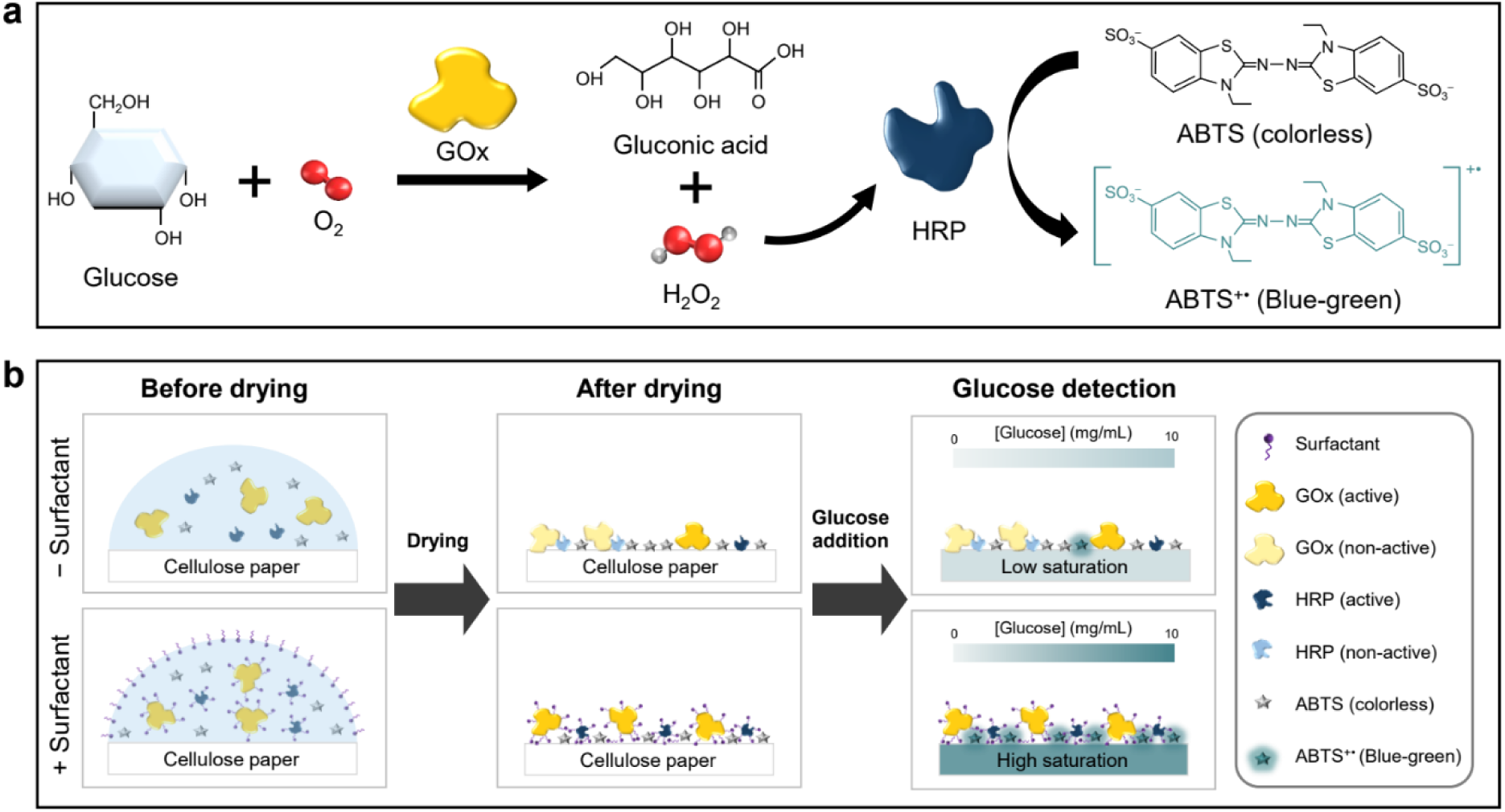
Principle of the surfactant-assisted colorimetric glucose assay. (a) Enzymatic cascade for glucose detection with GOx, HRP, and ABTS. (b) Schematic comparison of enzyme coating on cellulose paper without and with surfactant, shown in three steps: before drying, after drying, and glucose detection.

### 2.2 Screening and Optimization of Surfactants for Enhanced Detection Performance

To identify the most effective surfactant, comparative experiments were performed using nonionic surfactants (Tween 20, Tween 80, and Triton X-100), an anionic surfactant (SDS), and a multi-surfactant dish detergent. Each surfactant was mixed with GOx/HRP/ABTS, and the resulting mixture was coated onto paper and subjected to the glucose assay (Figure 2a, b). The colorimetric response was quantified using the chroma value (*C\**) in the CIE L*a*b* color space. Notably, under most tested conditions, 10 mg/mL glucose produced a lower chroma value than 5 mg/mL glucose. A supplementary analysis of dry-state absorbance at 780 nm similarly showed that, at high glucose concentrations, the absorbance no longer increased with increasing glucose concentration and instead decreased (Figure S3). Several factors may contribute to this signal decrease, including oxygen limitation, inhibition of enzymatic activity by excess H_2_O_2_, and instability of oxidized ABTS.^35–38^ However, the exact mechanism remains unresolved. In the *C\** measurements, the high-concentration signal decrease was observed in the control and most surfactant-treated conditions, suggesting that surfactant addition generally did not prevent this behavior. Accordingly, the analytical range was restricted to 0–5 mg/mL, over which *C\** increased monotonically, and the 10 mg/mL data point was excluded from curve fitting.

**Figure 2.**
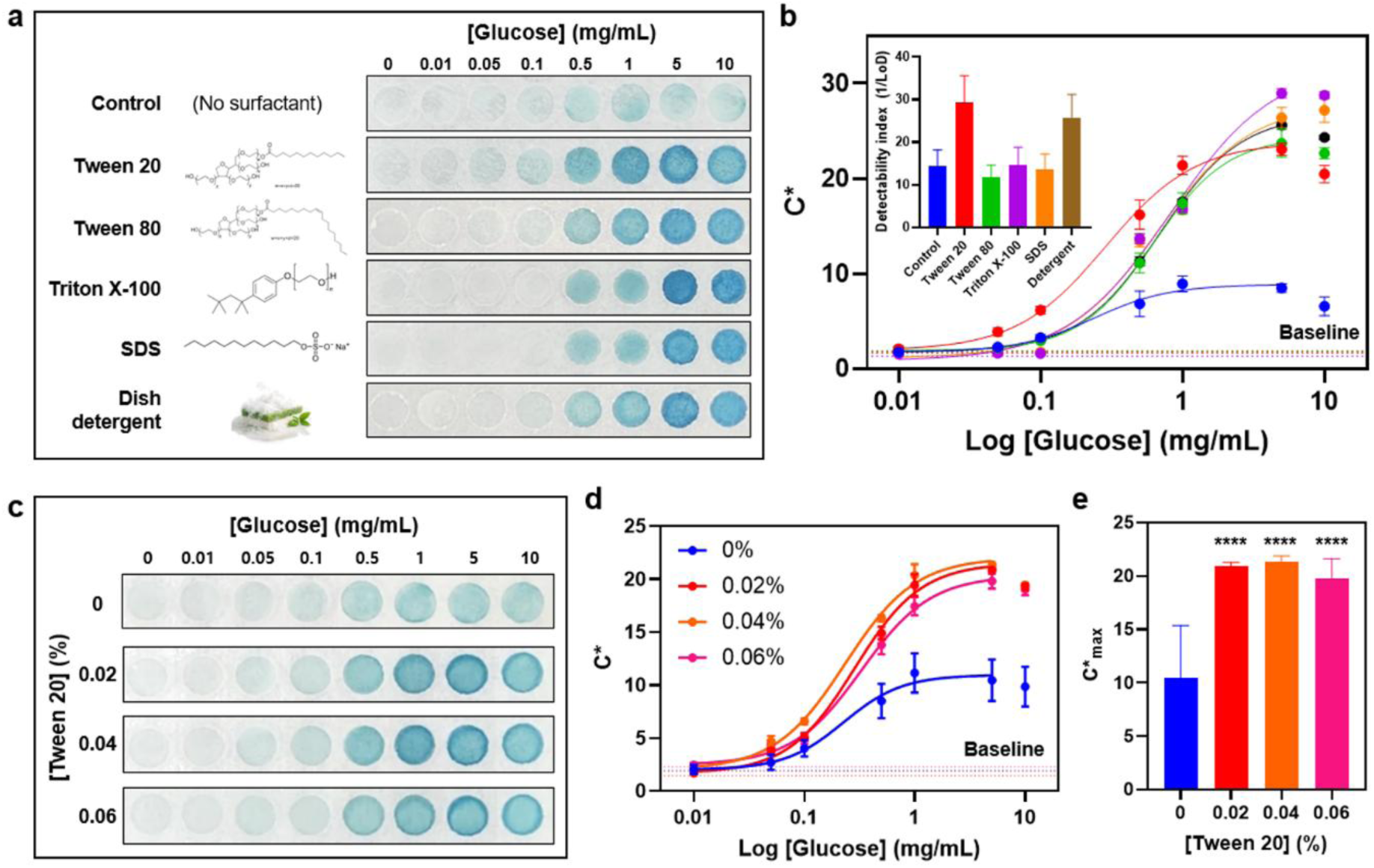
Effect of surfactants and optimization of Tween 20 concentration for the colorimetric glucose assay. (a) Representative photographs of the colorimetric glucose assay on GOx/HRP/ABTS-coated paper prepared without surfactant or with each surfactant; the chemical structure or image of each surfactant is shown at left. (b) Calibration curves of the chroma value (*C\**). Inset: detectability index ± SE, calculated as the reciprocal of the limit of detection (1/LoD) for each condition. (c) Representative photographs, (d) calibration curves, and (e) maximum chroma (*C\**_max_) for sensors prepared at increasing Tween 20 concentrations; \*\*\*\**p* < 0.0001 compared with 0% (v/v) Tween 20. Dashed lines in (b) and (d) indicate the baseline signal (*C\**at 0 mg/mL glucose) for each condition. Data are presented as mean ± SD (*n* = 3).

All surfactant-treated conditions exhibited markedly higher colorimetric responses than the control. The four-parameter logistic (4PL)-fitted colorimetric amplitude (span) was also greater than that of the control under all surfactant conditions, with Triton X-100 producing the largest span (Table S1). However, a different trend emerged when performance was evaluated using the detectability index, defined as the reciprocal of the limit of detection (1/LoD). Tween 20 and the dish detergent increased the detectability index by approximately 2.0- and 1.8-fold relative to the control, respectively. In contrast, the detectability indices obtained with Tween 80, Triton X-100, and SDS were approximately 0.8, 1.0, and 0.9 times that of the control, respectively, indicating no improvement in LoD. Thus, an increase in the absolute colorimetric response did not necessarily translate into improved detection performance. Although Tween 20 produced the greatest improvement in the detectability index, the relatively strong performance of the dish detergent was also noteworthy. Commercial dish detergents typically contain mixtures of multiple surfactants and other formulation components. Therefore, the relatively strong performance of the tested dish detergent raises the possibility of synergistic effects among its components. Although such effects were not directly examined in this study, this possibility represents an open question warranting further investigation using defined surfactant mixtures. Nevertheless, because the exact composition and concentrations of the individual components in the dish detergent could not be confirmed, its use presented limitations for assay standardization and mechanistic interpretation. Tween 20, which exhibited the highest detection performance among the individual surfactants with a well-defined composition, was therefore selected for subsequent experiments.

Having selected Tween 20 as the best-performing chemically defined surfactant, we then evaluated its concentration dependence over the range examined (Figure 2c–e). Increasing the Tween 20 concentration from 0.02 to 0.06% (v/v) produced no statistically significant differences in the maximum chroma value (*C\**_max_) among the three concentrations, while the LoD remained comparable across the tested concentrations (Table S2). Accordingly, 0.02% (v/v), the lowest concentration tested that produced the observed enhancement, was selected for subsequent experiments.

### 2.3 Tween 20-Enhanced Detection Performance under Low GOx Loading

In the preceding experiments, Tween 20 improved both the colorimetric response and detection performance. Because enzymes account for a substantial portion of the fabrication cost of paper-based enzymatic sensors, maintaining analytical performance while reducing GOx usage may provide a practical advantage by lowering reagent costs. To evaluate how the effect of Tween 20 varied with GOx loading, the GOx concentration was adjusted to 0.1, 1, and 10 mg/mL, and the effect of Tween 20 addition was compared at each concentration.

First, to determine whether the Tween 20-assisted enhancement in colorimetric response was related to changes in substrate wettability, the water contact angle was measured at each GOx concentration (Figure 3a, b). In the absence of Tween 20, the contact angle increased progressively with GOx concentration, from 25.78 ± 2.64° at 0.1 mg/mL to 42.20 ± 6.95° at 1 mg/mL and 66.93 ± 6.49° at 10 mg/mL. This trend suggests that the deposited enzyme increasingly influences the wetting behavior of the substrate at higher concentrations. Consistent with this interpretation, quantitative scanning electron microscopy (SEM) image analysis of the fiber network showed that the apparent surface porosity was significantly lower at 10 mg/mL GOx than in bare paper, whereas no significant difference from bare paper was detected at 0.1 mg/mL (Figure S4). This loading-dependent decrease in surface porosity is consistent with denser coverage of the cellulose fibers at high enzyme loading. Tween 20 significantly reduced the contact angle to 26.57 ± 1.53° at 1 mg/mL and 33.54 ± 5.27° at 10 mg/mL. At 0.1 mg/mL GOx, the mean contact angles were 25.78 ± 2.64° without Tween 20 and 27.11 ± 2.87° with Tween 20, and this difference did not reach statistical significance. Protein drying can increase the exposure of hydrophobic regions at the surface, and nonionic surfactants can associate directly with protein surfaces in addition to adsorbing at interfaces.^39^ Molecular dynamics simulations indicate that this association occurs at nonspecific regions of the protein surface, predominantly through the polyoxyethylene head groups.^40^ Such interactions could alter the hydration and apparent wettability of an enzyme-coated surface, with their effect becoming more readily detectable as protein coverage increases. At 0.1 mg/mL GOx, the absence of a detectable change in surface porosity relative to bare paper suggests that the hydrophilic cellulose network remained the dominant determinant of wetting behavior. Consequently, the surfactant-free substrate already exhibited a low contact angle, leaving limited scope for a further reduction.

**Figure 3.**
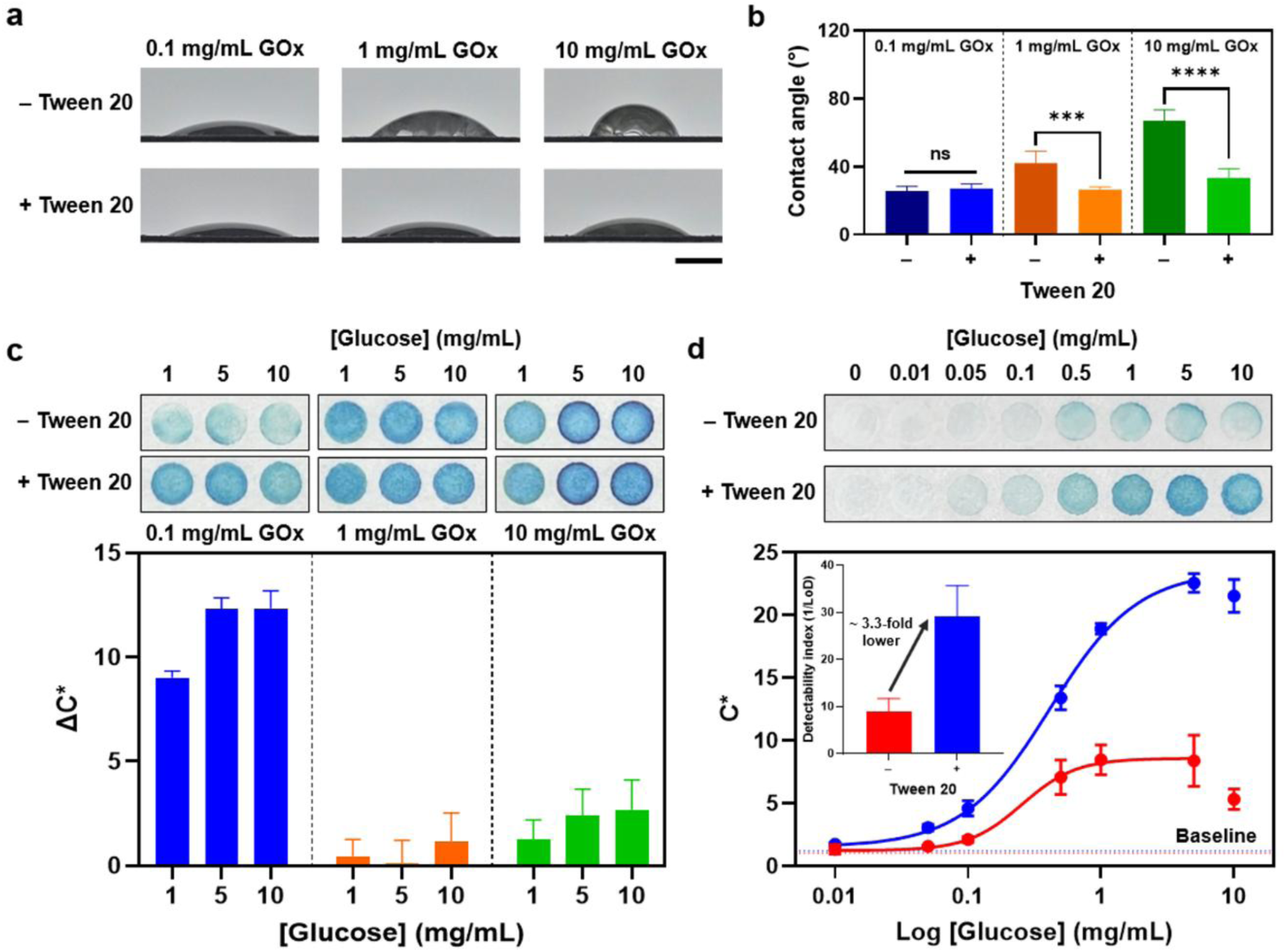
Water contact angle and colorimetric response of the glucose assay at different GOx concentrations with and without Tween 20. (a) Representative images of water droplets on sensors prepared with 0.1, 1, and 10 mg/mL GOx, without and with Tween 20 (scale bar: 2 mm), and (b) corresponding contact angles. *\*\*\*\*p* < 0.0001, *\*\*\*p* < 0.001; ns, not significant. (c) Representative photographs at selected glucose concentrations for each GOx concentration and the difference in *C\** between sensors without and with Tween 20 (*ΔC\**). (d) Representative photographs and calibration curves of the glucose assay at 0.1 mg/mL GOx. Inset: detectability index (1/LoD) ± SE without and with Tween 20. Dashed lines in (d) indicate the baseline signal. Data are presented as mean ± SD (n = 3).

The glucose-dependent colorimetric responses obtained with and without Tween 20 were then compared across the GOx concentrations (Figure 3c). Representative images further showed that Tween 20 produced visibly stronger color development at 0.1 mg/mL GOx. The increase in colorimetric response resulting from Tween 20 addition (Δ*C\**) was calculated at each glucose concentration. Tween 20 produced relatively small changes in colorimetric response at GOx concentrations of 1 and 10 mg/mL but markedly increased the signal at 0.1 mg/mL. At this low enzyme concentration, however, no statistically significant difference in contact angle was detected between the conditions with and without Tween 20. Although this nonsignificant result should not be interpreted as evidence of equivalent wettability, the pattern of the measured contact-angle changes did not parallel that of the signal enhancement. Tween 20 significantly reduced the contact angle at 1 and 10 mg/mL GOx, where the changes in colorimetric response were relatively small, whereas the most pronounced signal enhancement occurred at 0.1 mg/mL GOx. Therefore, the Tween 20-assisted enhancement of the colorimetric response cannot be fully explained by the measured macroscopic wettability alone.

At 0.1 mg/mL GOx, which exhibited the largest Tween 20 effect, the colorimetric response was further examined across the full glucose concentration range (Figure 3d). Analysis of the resulting calibration curve showed that Tween 20 decreased the LoD from 0.113 ± 0.036 to 0.034 ± 0.008 mg/mL (Table S3), corresponding to an approximately 3.3-fold increase in the detectability index (Figure 3d, inset). The pronounced reduction in the LoD at 0.1 mg/mL GOx further highlights the mismatch between signal enhancement and the loading-dependent changes in wettability described above, as it occurred under the low-loading condition in which no statistically significant contact-angle difference was detected with Tween 20 and the apparent surface porosity did not differ significantly from that of bare paper. Taken together, these findings suggest that the enhancement cannot be attributed primarily to changes in macroscopic wettability or dense coverage of the cellulose fibers and point to additional effects within the dried enzyme formulation. For clinical context, the assay exhibited a monotonic analytical range of 0–5 mg/mL (0–500 mg/dL), covering the concentration threshold for pathologic glucosuria (>25 mg/dL) and the detection thresholds of conventional semiquantitative urine glucose tests (50–250 mg/dL).^41^ Although this range supports the potential application of the assay to urinary glucose screening, validation using clinical urine samples remains necessary.

### 2.4 Effect of Tween 20 on Enzyme Distribution in Solution and the Dried State

The Tween 20-induced performance enhancement was not fully explained by changes in substrate wettability (Figure 3). We therefore examined the effect of Tween 20 on enzyme distribution and aggregation in both solution and the dried state. First, dynamic light scattering (DLS) was used to determine whether Tween 20 affected enzyme aggregation in solution (Figure S5). Addition of Tween 20 to the GOx/HRP sample produced no notable change in the position of the primary peak or the overall size distribution, indicating that any change in enzyme aggregation detectable by DLS in solution was minimal.

To examine interfacial effects and changes in enzyme distribution that may arise during drying, the dried coatings were first analyzed by SEM at micrometer scale (Figure S4). The SEM images primarily revealed the underlying cellulose-fiber network, and no consistent Tween 20-dependent change in the overall fiber architecture was apparent. However, SEM was not sufficient to reliably resolve the nanoscale enzyme layer or to distinguish protein-derived features from the cellulose substrate and other dried formulation components. Nanoscale AFM analysis directly on paper was also difficult because the high surface roughness of paper complicates reliable nanoscale morphological interpretation. Therefore, the same enzyme mixtures were coated onto a flat mica substrate, dried, and examined by AFM (Figure 4). Freshly cleaved mica was selected as a model substrate because it provides an atomically flat, hydrophilic reference surface on which protein adsorption has been extensively characterized. It was therefore used to visualize how Tween 20 alters the organization of the dried enzyme layer rather than to reproduce the cellulose surface itself. However, because mica and cellulose differ in surface charge, chemical composition, roughness, and porosity, the morphology observed on mica may be substrate-dependent and should not be interpreted as a direct representation of the enzyme distribution on paper. AFM-derived topographical parameters, including height and lateral dimensions, can provide a basis for distinguishing deposited nanoscale structures with different morphologies.^42–45^ The GOx/HRP coating exhibited a continuous, laterally spread surface morphology without clearly distinguishable individual structures (Figure 4a, b). In contrast, the GOx/HRP/Tween 20 coating exhibited protruding structures that were relatively dispersed across the surface (Figure 4c, d). To further compare the observed differences in height distribution, the surface skewness (*S*_sk_) was analyzed (Figure S6e). *S*_sk_ quantifies the asymmetry of the height distribution, with larger positive values indicating a greater contribution of elevated particles to the height distribution.^46, 47^ The *S*_sk_ of the GOx/HRP coating was −0.0803 ± 0.1614, indicating a nearly symmetric height distribution, whereas that of the GOx/HRP/Tween 20 coating was 0.4893 ± 0.1536, indicating a positively skewed height distribution. This increase in *S*_sk_ was consistent with the protruding structures observed in the AFM images. AFM analysis of individually coated GOx and HRP layers further confirmed that the two enzymes exhibited distinct surface morphologies after drying, providing additional context for interpreting the mixed GOx/HRP coating (Figure S6a–d). Additional ImageJ analysis showed a greater number of elevated particles and a larger average area per detected particle in the Tween 20-containing coating than in the GOx/HRP coating (Figure S6f, g). Together, these results were consistent with a more discrete and feature-rich dried-film morphology following Tween 20 addition rather than a predominantly continuous deposit.

**Figure 4.**
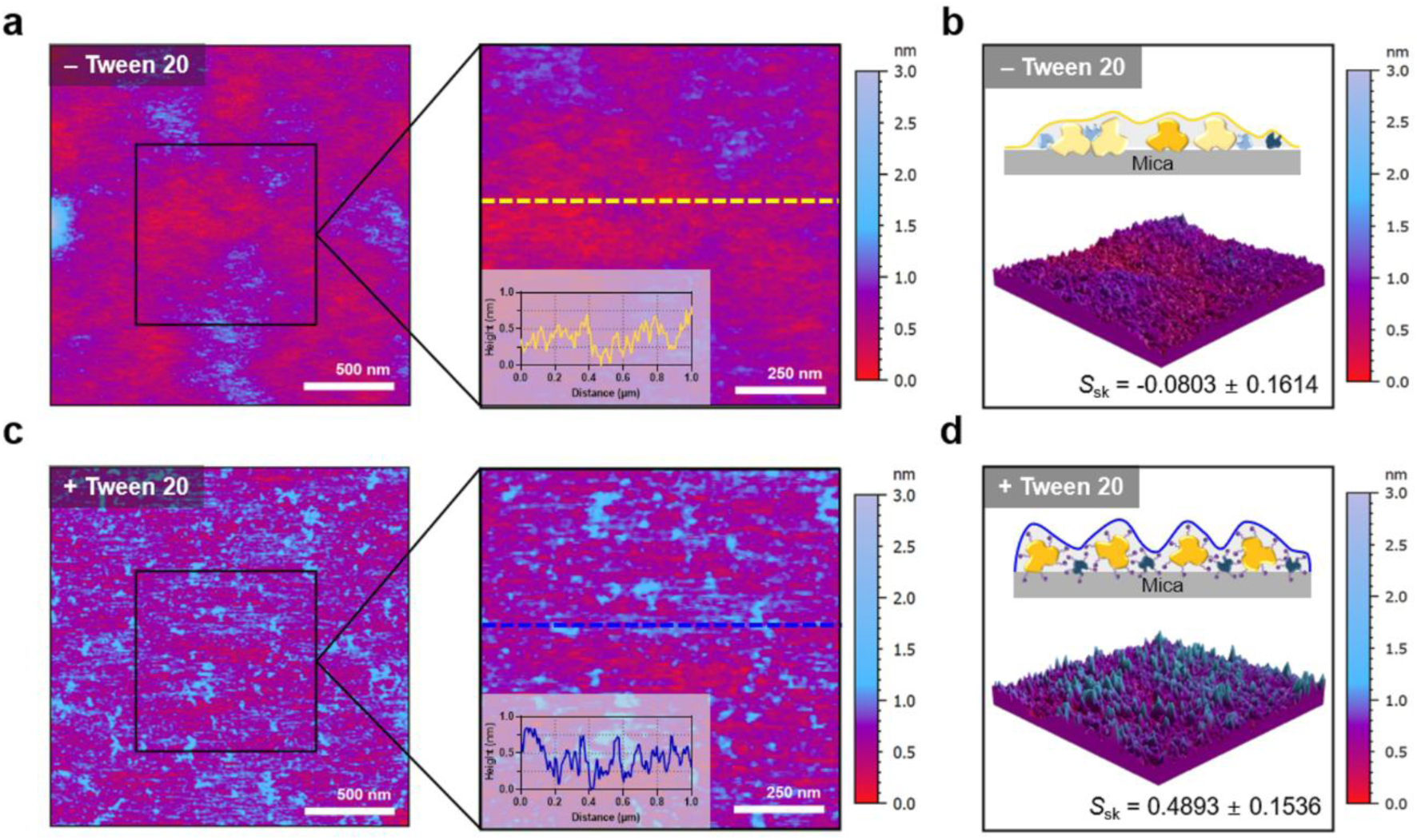
AFM analysis of enzyme-coated mica with and without Tween 20. (a) Height image of GOx/HRP-coated mica (–Tween 20, 2 × 2 μm^2^, left) with a magnified image (1 × 1 μm^2^, right); the inset shows the cross-sectional height profile along the dashed line in the magnified image. (b) 3D topographic image of the same surface, with a schematic illustration of the enzyme arrangement on mica. (c, d) Corresponding images for GOx/HRP/Tween 20-coated mica (+Tween 20). *S*_sk_: surface skewness. Scale bars: 500 nm (full images), 250 nm (magnified images).

These observations are consistent with the reported protein-stabilizing actions of surfactants. Nonionic surfactants can preferentially adsorb at the air–water interface, thereby reducing interfacial exposure and adsorption of proteins,^23, 32^ or interact with hydrophobic regions on protein surfaces to limit protein aggregation.^48^ Consistent with this protective role, polysorbate 20 (PS20) was reported to markedly reduce subvisible particle formation in lyophilized immunoglobulin formulations containing trehalose; addition of PS20 to the reconstitution medium also reduced particle formation.^49^ The molecular dynamics simulations noted above further indicate that association of Tween 20 with the protein surface reduces the conformational space sampled by the protein.^40^ Although these simulations were performed with bovine serum albumin rather than GOx or HRP, they provide a mechanistic precedent for the possibility that similar surfactant–protein interactions may contribute to the enzyme stabilization observed in our system. Because enzymes encounter both air–water and solid–water interfaces during coating and drying, Tween 20 may mitigate interfacial adsorption and intermolecular aggregation through these pathways. This interpretation is consistent with the AFM results, which showed a more dispersed surface distribution of the dried enzymes in the presence of Tween 20. Because excessive enzyme aggregation can promote conformational changes and activity loss, this altered surface distribution may have contributed to the preservation of enzyme cascade activity after drying. Although the AFM observations do not establish the molecular mechanism of enzyme stabilization, they provide morphological evidence supporting the proposed protective effect of Tween 20 during drying.

The stronger effect of Tween 20 at low GOx loading may reflect the greater proportional impact of interfacial enzyme loss under enzyme-limited conditions. Because the sample volume and reaction area were fixed, the available interfacial area remained comparable across the tested GOx concentrations. If a similar absolute amount of enzyme was lost through interfacial adsorption at each loading, this loss would account for a larger fraction of the total enzyme at lower GOx concentrations. At higher GOx loading, sufficient active enzyme may remain after drying, thereby reducing the difference between the conditions with and without Tween 20. To test this proposed protective effect directly, the time-dependent colorimetric responses were compared in solution phase and on paper before and after a controlled dry–rehydrate cycle (Figure S7). In solution, the responses obtained with and without Tween 20 were comparable, indicating that Tween 20 does not itself alter the catalytic behavior of the cascade in the absence of a drying step. After drying, the colorimetric response across the glucose concentrations decreased markedly in the absence of Tween 20, whereas Tween 20 attenuated this decrease, although the response remained below that observed before drying. The same trend was evident in the reaction kinetics: drying reduced the apparent rate constant from 0.0143 to 0.0015 s^-1^ without Tween 20 (11% retention), compared with 0.0130 to 0.0123 s^-1^ with Tween 20 (95% retention). Because the paper-deposited conditions differed only in whether the reagent was dried, the contribution of the substrate was common to both, and the comparison therefore isolated the effect of drying itself. Having established that Tween 20 preserves cascade activity across a dry–rehydrate cycle, we next examined whether this is reflected in the analytical performance of sensors subjected to prolonged stress.

### 2.5 Effect of Tween 20 on Sensor Stability and Selectivity

To evaluate whether Tween 20 improved the retention of the analytical response under stress conditions, the sensors were subjected to three stress treatments. Wetting–drying and thermal stresses were applied to the sensors before glucose treatment, whereas ionic strength stress was evaluated during glucose detection using glucose solutions containing NaCl. Changes in colorimetric intensity with repeated wetting–drying cycles and thermal stress duration at 90 °C were fitted separately using a one-phase decay model, and the resulting decay rate constants (*k*) were compared. The *k* value of the Tween 20-treated sensor was approximately 1.7-fold lower than that of the control under repeated wetting–drying cycles (Figure 5a) and 2.2-fold lower under thermal stress (Figure 5b). A lower *k* value indicates a slower decline in the analytical signal with increasing stress exposure. Accordingly, the Tween 20-treated sensors retained their glucose assay performance more effectively under repeated moisture exposure and elevated temperature.

**Figure 5.**
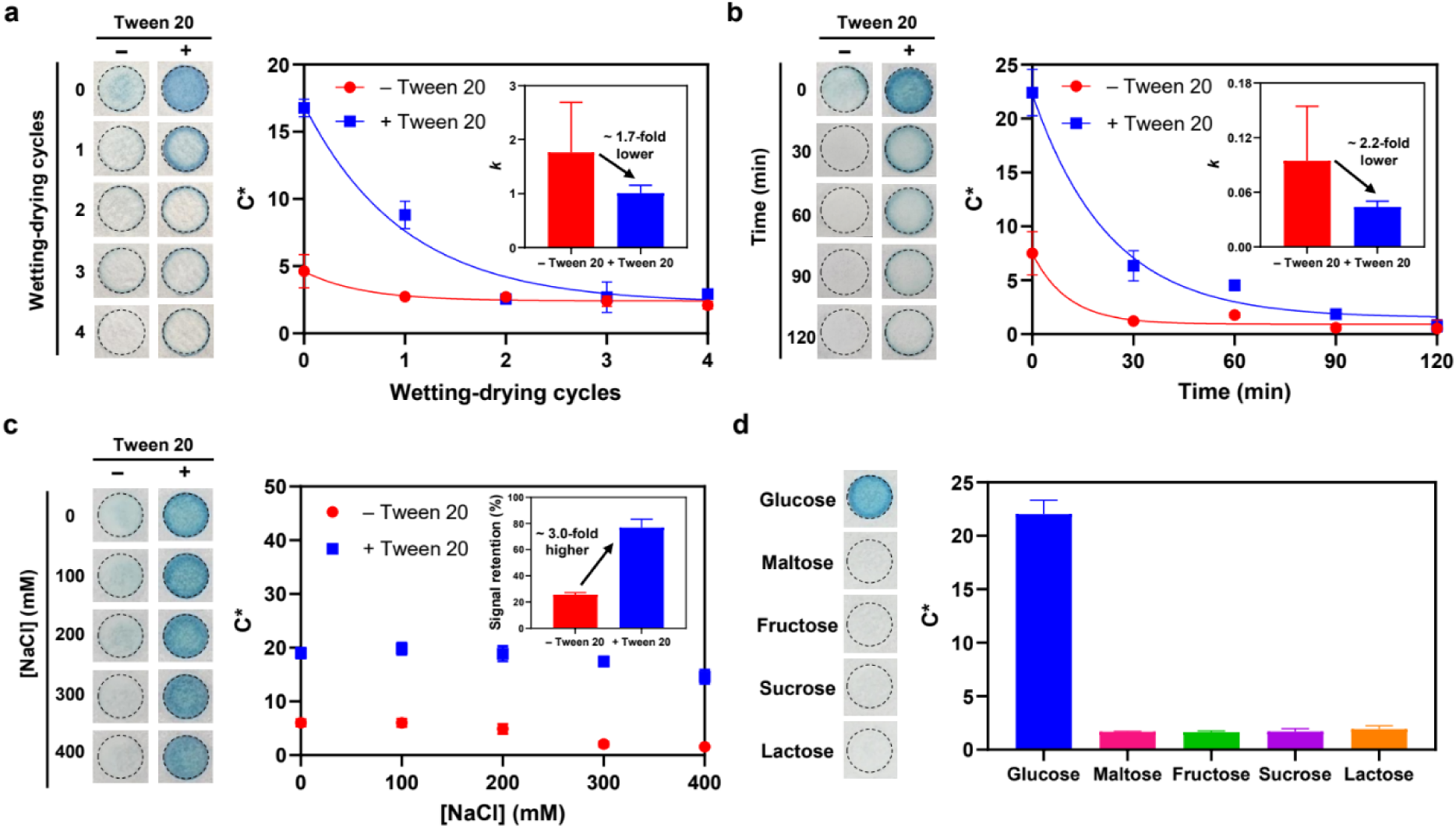
Stability under stress conditions and selectivity of the glucose sensor. (a–c) Representative photographs and *C\** of sensors prepared without and with Tween 20 under stress conditions: (a) repeated wetting–drying cycles, (b) heating at 90 °C, and (c) increasing NaCl concentration. Insets: decay rate constant (*k*) ± SE in (a) and (b), and signal retention at 400 mM relative to 0 mM in (c). (d) Representative photographs and *C\** of the assay for glucose and other sugars. Data are presented as mean ± SD (*n* = 3).

Under the ionic strength stress condition, the decay curve did not approach a sufficiently defined plateau within the evaluated NaCl concentration range, precluding reliable estimation of *k*. Instead, relative signal retention was calculated by comparing the colorimetric response at the highest concentration tested, 400 mM NaCl, with that at 0 mM NaCl. The Tween 20-treated sensor retained approximately 77% of the colorimetric response observed at 0 mM NaCl, whereas the sensor without Tween 20 retained only approximately 26% (Figure 5c). Thus, Tween 20 markedly suppressed the loss of the colorimetric response under high ionic strength conditions.

The slower signal loss observed following repeated wetting–drying cycles and thermal stress, together with the greater signal retention during glucose detection under high ionic strength conditions, supports the interpretation that Tween 20 helps preserve enzymatic performance. These results are also consistent with the more dispersed surface distribution observed by AFM, suggesting that reduced aggregation may contribute to the retention of enzymatic performance during drying and stress exposure. Nevertheless, the final colorimetric signal may be influenced not only by enzyme activity but also by ABTS stability, reconstitution of the dried reagents, and mass transport within the substrate.

Because surfactant–enzyme interactions could potentially alter enzyme structure or substrate recognition, we further examined whether Tween 20 affected the selectivity of the GOx-based assay. In addition, because real samples typically contain sugars other than glucose, it was necessary to verify that the sensor retained glucose selectivity in the presence of the surfactant. To this end, the colorimetric responses to potentially interfering sugars were compared (Figure 5d). A distinct colorimetric response was observed only for glucose, with no significant response detected for the other sugars. This result indicates that the substrate selectivity of the GOx-based reaction for glucose was maintained in the presence of Tween 20.

Additional interference, matrix, and reproducibility studies were conducted to evaluate the analytical robustness of the assay. Tween 20 attenuated the effects of uric acid, ascorbic acid, and acetaminophen, although cysteine still caused a modest reduction in the colorimetric response (Figure S8). In artificial saliva, Tween 20 decreased the LoD from 0.186 ± 0.076 mg/mL to 0.092 ± 0.025 mg/mL, demonstrating that the detection benefit was retained in the presence of matrix components, although both LoDs were higher than those obtained in DW (Figure S9). Reproducibility was also improved: all within-PPP and between-batch coefficients of variation (CVs) were below 10% with Tween 20, whereas the corresponding values without Tween 20 ranged from 12.3 to 19.5% (Figure S10 and Table S4). To determine whether the enhancement was specific to ABTS, the assay was further evaluated using 3,3′,5,5′-tetramethylbenzidine (TMB) as an alternative chromogen (Figure S11). Tween 20 similarly improved detectability, indicating that the effect was not restricted to a single chromogenic substrate. However, the formation of dark precipitates at high glucose concentrations limited quantitative analysis to the lower concentration range. Together, these additional analyses indicate that the beneficial effect of Tween 20 extends beyond the aqueous ABTS assay, while also identifying residual matrix- and chromogen-dependent limitations. Key quantitative comparisons are summarized in Table S5.

## 3. Conclusion

In this study, we demonstrated a surfactant-assisted strategy for improving the colorimetric response and detection performance of a paper-based glucose assay under reduced enzyme loading. Although all tested surfactants increased colorimetric intensity, only selected surfactants improved the LoD, with Tween 20 providing the greatest enhancement among the chemically defined surfactants. A Tween 20 concentration of 0.02% (v/v) was identified as the lowest concentration tested that produced the observed enhancement, and its effect became more pronounced as GOx loading decreased, lowering the LoD from 0.113 to 0.034 mg/mL at 0.1 mg/mL GOx. This improvement occurred without a significant change in contact angle, indicating that the effect of Tween 20 cannot be fully explained by the measured macroscopic wettability alone. Tween 20 did not appreciably alter the enzyme size distribution or reaction kinetics in solution, but it preserved the apparent rate constant of the GOx–HRP cascade after drying, and AFM revealed a more dispersed dried-state enzyme distribution. Together with improved signal retention under wetting–drying, thermal, and high-ionic-strength conditions, improved reproducibility, and maintained glucose selectivity, these results indicate that Tween 20 helps retain enzyme cascade activity during drying. Enzyme-specific activity measurements and quantification of interfacial adsorption will be required to identify the contributions of GOx and HRP and the underlying mechanism. Overall, the surfactant-assisted 96-PPP provides a simple, parallelizable platform for systematically distinguishing signal enhancement from genuine improvements in detection performance across enzyme-loading and formulation conditions. By improving assay response and robustness at reduced GOx loading without additional immobilization chemistry, this strategy offers a practical route toward lower reagent consumption, more efficient formulation optimization, and enhanced performance of dried paper-based enzymatic sensors.

## 4. Materials & Methods

### Materials

Glucose oxidase from *Aspergillus niger* (GOx), horseradish peroxidase (HRP), 2,2’-azino-bis(3-ethylbenzothiazoline-6-sulfonic acid) diammonium salt (ABTS), Whatman 41 filter paper, Tween 20, Tween 80, Triton X-100, sodium chloride (NaCl), D-(+)-glucose, D-(-)-fructose, D-(+)-maltose monohydrate, sucrose, D-lactose monohydrate, uric acid, L-ascorbic acid, L-cysteine, potassium chloride, potassium phosphate monobasic, urea, ammonium chloride, calcium chloride dihydrate, sodium bicarbonate, sodium sulfate, and TMB were purchased from Sigma-Aldrich (USA). A 10% (w/v) SDS solution was purchased from Biosesang (South Korea), dish detergent and Tylenol (acetaminophen) were purchased from a local grocery store in South Korea. Distilled water (DW) was purchased from Gibco (USA).

### Fabrication of the 96-PPP Glucose Sensor

The paper-based sensor platform, 96-PPP, was prepared by patterning hydrophilic puddles on Whatman 41 filter paper, following a previously reported fabrication method.^33^ To prepare the reagent solution, GOx (20 mg/mL), HRP (5 mg/mL), ABTS (20 mM), and Tween 20 (1% (v/v)) were diluted with 150 mM phosphate–citrate buffer to final concentrations of 0.1 mg/mL GOx, 5 μg/mL HRP, 5 mM ABTS, and 0.02% (v/v) Tween 20. Subsequently, 10 μL of the reagent solution was deposited onto each puddle of the 96-PPP and dried in a food dehydrator (Shinil, South Korea) at approximately 35 °C for 30 min. The dehydrator operates by forced-air convection without active humidity control; the ambient relative humidity during fabrication was 20–40%. The sensors were used after complete drying.

### 96-PPP-Based Glucose Assay

Glucose solutions were prepared at various concentrations (0.00, 0.01, 0.05, 0.10, 0.50, 1.00, 5.00, and 10.00 mg/mL) to evaluate the analytical performance of the paper-based glucose sensor. For each concentration, 5 μL of glucose solution was dropped onto the sensors, and the color response of the sensor was monitored for 5 min, and the image acquired at 5 min was used for quantification. Images were acquired using a smartphone camera (Galaxy S23, Samsung, South Korea) mounted on a fixed stand at a constant distance and angle under ambient fluorescent illumination, with white balance and exposure left at the camera’s automatic settings. All conditions to be compared within a given experiment were recorded within a single frame.

### Optimization of Reagent Deposition and Buffer Conditions

The effect of sequential reagent deposition on the colorimetric response of the paper-based glucose sensor was examined while maintaining the same final reagent composition. Individual reagent solutions were deposited onto the paper-based platform in different sequences, with each deposition step followed by drying. The final concentrations of GOx, HRP, ABTS, and Tween 20 were maintained at 0.1 mg/mL, 5 μg/mL, 5 mM, and 0.02% (v/v), respectively. After completing the sequential deposition process, 5 μL of glucose solution was applied to each sensor, and the resulting colorimetric response was monitored and compared.

The effect of buffer conditions on the colorimetric response of the paper-based glucose sensor was also examined. The reagent solutions were prepared using DW, 20 mM phosphate buffer, or 150 mM phosphate–citrate buffer while maintaining the same final concentrations of GOx, HRP, ABTS, and Tween 20. After fabrication of the sensors, 10 μL of glucose solution was applied to each sensor, and the resulting colorimetric response was monitored and compared.

### Dry-State Absorbance Measurement

To examine the colorimetric response at high glucose concentrations, 10 μL of glucose solution (0–10 mg/mL) was applied to GOx/HRP/ABTS-coated sensors. The sensors were then dried at room temperature for 2 h, and the absorbance of each reaction puddle at 780 nm was measured using a microplate reader (Synergy H1, BioTek, USA).

### Surfactant Screening and Tween 20 Concentration Optimization

To compare the effects of different surfactants on the paper-based glucose sensor, Tween 20 in the reagent solution was replaced with Tween 80, Triton X-100, SDS, or dish detergent. All surfactants were evaluated under the same experimental conditions, and the final concentration of each surfactant was adjusted to 0.02% (v/v). The concentrations of GOx, HRP, and ABTS were maintained at the same values used in the preceding experiments. To optimize the Tween 20 concentration, sensors were prepared with final Tween 20 concentrations of 0, 0.02, 0.04, and 0.06% (v/v), while the concentrations of GOx, HRP, and ABTS were kept constant. The glucose assay was subsequently performed under the conditions described above, and the resulting colorimetric responses were compared.

### Evaluation under Different GOx Loading Conditions

To investigate the effect of Tween 20 at different GOx concentrations, paper-based glucose sensors were prepared using GOx concentrations of 10, 1, and 0.1 mg/mL. Except for the GOx concentration, all other experimental conditions, including the concentrations of HRP, ABTS, and Tween 20, were kept constant. After drying, 5 μL of glucose solution at concentrations of 1, 5, and 10 mg/mL was applied to each sensor. The colorimetric responses were monitored and compared to evaluate the dependence of the Tween 20 effect on the GOx concentration. To further evaluate detection performance under the low GOx loading condition, sensors prepared with 0.1 mg/mL GOx, with and without Tween 20, were tested using glucose concentrations of 0.00, 0.01, 0.05, 0.10, 0.50, 1.00, 5.00, and 10.00 mg/mL.

### Water Contact Angle Measurements

The wettability of the sensors prepared with different GOx concentrations was evaluated by water contact angle measurements. Sensors were fabricated using GOx concentrations of 10, 1, and 0.1 mg/mL, while other reagent conditions were kept constant. A 20 μL droplet of DW was placed onto the surface of each sensor, and images of the droplet were acquired from the side using a Galaxy S23. The contact angle was measured from the acquired images using ImageJ (NIH, USA).

### Scanning Electron Microscopy (SEM) Analysis

The surface morphology of the paper substrates was examined using SEM (CX-200K, COXEM, South Korea) at an accelerating voltage of 15 kV. Untreated paper was used as the bare paper control. For reagent-treated samples, the reagent solution was prepared under the same conditions as those used in the colorimetric assay, with the GOx concentration set to 0.1 or 10 mg/mL, each without and with Tween 20. Each solution was applied to the paper substrate and dried under the same conditions used for the paper-based assay. Prior to imaging, the samples were sputter-coated with Au for 240 s at 5 mA using an ion sputter coater (SPT-20, COXEM, South Korea). Images were acquired at 500×, 1000×, and 3000×. For porosity quantification, 500× images were binarized in ImageJ using the Otsu auto-threshold method, and the pore area fraction was determined using the Analyze Particles function.

### Dynamic Light Scattering (DLS) Analysis

The hydrodynamic size distribution of the enzyme samples was measured by DLS using a Zetasizer Nano ZS (ZEN3600, Malvern, UK) at 25 °C. GOx/HRP mixtures were prepared with or without Tween 20, and their particle size distributions were analyzed. The final concentrations of GOx, HRP, and Tween 20 were maintained at 0.1 mg/mL, 5 μg/mL, and 0.02% (v/v), respectively, consistent with the conditions used for the paper-based glucose sensor. Each size distribution represents the average of three repeated measurements of the same sample.

### Atomic Force Microscopy (AFM) Characterization

The morphology of the enzyme samples was analyzed using AFM (MultiMode 8, Bruker, USA). GOx and HRP were analyzed separately, and a GOx/HRP mixture was also prepared. To examine the effect of Tween 20 on enzyme morphology, a GOx/HRP mixture containing Tween 20 was additionally prepared. The final concentrations of GOx, HRP, and Tween 20 were adjusted to 0.1 mg/mL, 5 μg/mL, and 0.02%, respectively. Freshly cleaved mica was used as a flat model substrate to facilitate nanoscale topographic imaging of the dried enzyme layer rather than to reproduce the cellulose-paper surface. Then, 10 μL of each sample solution was deposited onto freshly cleaved mica and incubated for 5 min. The mica surface was gently washed with DW to remove unbound molecules and dried under a stream of nitrogen gas. Images were acquired in tapping mode using PR-T300 cantilevers (Probes, South Korea) over scan areas of 2 × 2 μm^2^. Magnified images (1 × 1 μm^2^) were cropped from these images for further analysis. Height images were analyzed using MountainsSPIP (version 9, Digital Surf, France). *S*_sk_ was calculated from the height distribution of the cropped 1 × 1 μm^2^ regions (Equation 1).^46^ For each condition, three independent regions were analyzed, and *S*_sk_ values are reported as mean ± SD.

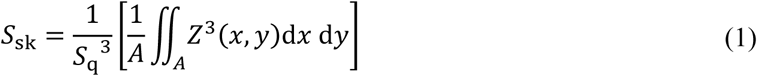

where *Z* (*x*, *y*) is the height at each point relative to the mean plane and A is the sampling area.

For particle analysis, height images displayed on an identical z-scale (0–3 nm) were converted to RGB format in ImageJ, and the green channel, which corresponded to the elevated regions in the color scale, was extracted. The resulting images were binarized using the Otsu auto-threshold method, and the number and area of particles were quantified using the Analyze Particles function with a minimum particle size of 50 nm^2^ to exclude noise.

### Retention of Enzyme Cascade Activity after Drying

To evaluate the effect of drying on the enzymatic cascade reaction, the colorimetric reaction was compared under three conditions, each without and with Tween 20: in solution, on paper without drying, and on paper after drying. The reagent composition was identical to that used in the paper-based assay. For the solution-phase condition, 40 μL of the reagent solution and 20 μL of glucose solution were mixed in a transparent 96-well plate. This maintained the reagent-to-sample volume ratio of the paper-based assay (10 μL of reagent and 5 μL of glucose solution) while ensuring complete coverage of the well bottom. For the undried paper condition, 10 μL of the reagent solution was dispensed onto each reaction puddle, and 5 μL of glucose solution was applied immediately. For the dried paper condition, the reagent solution was dispensed and dried as described above, and 5 μL of glucose solution was then applied to rehydrate the dried reagents and initiate the reaction. Colorimetric responses to 0–10 mg/mL glucose were recorded 5 min after glucose addition. For kinetic analysis, 5 mg/mL glucose was used, and images were acquired every 10 s for 290 s. The apparent rate constant was determined as described below.

### Assay Performance under Stress Conditions

Stability was evaluated under three stress conditions: repeated wetting–drying cycles, thermal stress, and ionic strength stress. For wetting–drying stress, 5 μL of DW was applied to each reaction puddle and dried in a food dehydrator at 35 °C for 30 min. This cycle was repeated 0, 1, 2, 3, or 4 times. For thermal stress, the sensors were placed in an oven at 90 °C for 0, 30, 60, 90, or 120 min. After each wetting–drying or thermal treatment, 5 μL of 5 mg/mL glucose solution was applied to each sensor, and the colorimetric response was monitored.

Ionic strength stress was evaluated during glucose detection using glucose solutions containing NaCl. The solutions were prepared with a constant glucose concentration of 5 mg/mL and NaCl concentrations of 0, 100, 200, 300, or 400 mM. A 5 μL aliquot of each solution was applied to the sensor, and the colorimetric response was monitored.

### Glucose Selectivity against Structurally Related Sugars

To evaluate the selectivity of the paper-based glucose sensor, the sensor was tested against glucose and structurally related sugars, including maltose, fructose, sucrose, and lactose. Each solution was prepared at a concentration of 5 mg/mL. A 5 μL aliquot of each solution was applied to the sensor, and the resulting colorimetric response was monitored and compared with that of glucose.

### Interference Test

To evaluate the interference resistance of the assay, the colorimetric response to 5 mg/mL glucose was measured in the absence and presence of potential interfering substances. Uric acid was dissolved in a 250 mM NaOH solution. Ascorbic acid and L-cysteine were dissolved directly in DW. Acetaminophen solutions were prepared by dissolving commercial Tylenol tablets (500 mg acetaminophen per tablet) in DW without removal of excipients. Uric acid, ascorbic acid, L-cysteine, or acetaminophen was individually added to the glucose solution at a final concentration of 0.1 mg/mL. A 5 μL aliquot of each solution was applied to sensors prepared without and with Tween 20, and the resulting colorimetric responses were compared with that of glucose alone.

### Glucose Detection in Artificial Saliva

Artificial saliva was prepared according to a previously reported composition by dissolving NaCl (0.0628 g), KCl (0.482 g), KH₂PO₄ (0.327 g), urea (0.100 g), Na₂SO₄ (0.382 g), NH₄Cl (0.0890 g), CaCl₂·2H₂O (0.114 g), and NaHCO₃ (0.315 g) in DW to a final volume of 500 mL, without pH adjustment, and was used shortly after preparation.^50^ Glucose was subsequently spiked into the prepared artificial saliva to obtain concentrations (0–10 mg/mL). A 5 μL aliquot of each sample was applied to sensors prepared without and with Tween 20. The LoD was determined over the 0–5 mg/mL range as described below and compared with that obtained in DW.

### TMB-based Colorimetric Assay

To determine whether the effect of Tween 20 was maintained with a different chromogenic substrate, ABTS was replaced with TMB. A 20 mM TMB stock solution was prepared in ethanol and diluted to a final concentration of 5 mM in the reagent solution. The concentrations of GOx, HRP, and Tween 20 and all other fabrication and assay procedures were identical to those used in the ABTS-based assay. Glucose solutions (0–10 mg/mL) were applied to sensors prepared without and with Tween 20.

### Image-Based Colorimetric Analysis

Images of the sensors were analyzed in the CIE L*a*b* color space using ImageJ. For each reaction puddle or well, the mean a* and b* values were extracted, and *C\** was calculated using Equation 2 and used as the colorimetric readout throughout this study.^51^

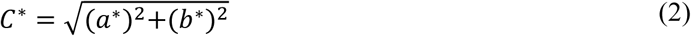

The difference in *C\** between sensors prepared with and without Tween 20 (Δ*C\**) was calculated by subtracting the *C\** of the sensor without Tween 20 from that of the sensor with Tween 20. Calibration curves of *C\** against glucose concentration were fitted with a four-parameter logistic (4PL) model (Equation 3) using GraphPad Prism 11.0.0 (GraphPad Software, USA).^52^

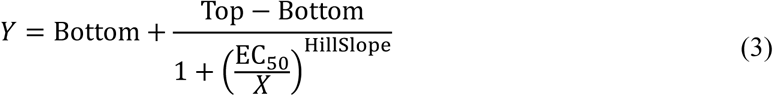

where *X* is the glucose concentration, *Y* is the chroma value, Bottom and Top are the lower and upper plateaus, respectively, EC_50_ is the glucose concentration corresponding to the midpoint between Bottom and Top, and HillSlope describes the steepness of the fitted curve. The span (Top − Bottom) was used as a measure of the amplitude of the colorimetric response.

The LoD was estimated by interpolating the 4PL curve at a response equal to the bottom plateau plus three times its standard error, following a previously reported method.^53^ The SE of the interpolated LoD was calculated from its 95% confidence interval as (upper limit − lower limit)/3.92, and the SE of the detectability index (1/LoD) was estimated by error propagation as SE(LoD)/LoD^2^. Curves were fitted using glucose concentrations in linear units (mg/mL); logarithmic axes were used only for display. Because *C\** decreased at 10 mg/mL relative to 5 mg/mL, fitting was restricted to 0–5 mg/mL for the ABTS-based assays. For the TMB-based assay, fitting was restricted to 0–1 mg/mL because dark precipitates formed at higher glucose concentrations.

For the wetting–drying and thermal stress tests, the decrease in *C\** was fitted with a one-phase exponential decay model (Equation 4), and the rate constant (*k*) was obtained from the fit.^52^

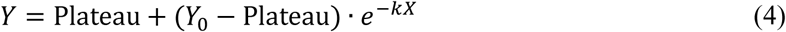

where *Y*_0_ is the initial response, Plateau is the asymptotic response, *X* is the number of wetting–drying cycles or the thermal stress duration, and *k* is the decay rate constant (cycle^-1^ or min^-1^).

For the ionic-strength test, the response did not reach a plateau within the tested NaCl range, and a decay constant could not be reliably estimated. Therefore, retention was calculated as the ratio of *C\** at 400 mM NaCl to that at 0 mM NaCl, expressed as a percentage.

For the drying experiments, the time-dependent increase in *C\** at 5 mg/mL glucose was fitted with a one-phase association model (Equation 5).^52^

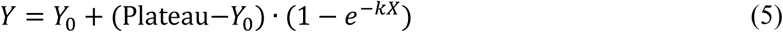

where *Y_0_* and Plateau are the initial and maximum responses, *X* is the reaction time and *k* is the apparent rate constant (s^-1^). Retention after drying was calculated as the ratio of *k* on dried paper to that on undried paper, expressed as a percentage.

### Statistical Analysis

Unless otherwise stated, *n* = 3 denotes three independently dispensed and dried reaction puddles on a single 96-PPP, prepared from one reagent solution; data are presented as mean ± standard deviation (SD). Parameters obtained from nonlinear regression (span, Hill slope, LoD, detectability index, and rate constants) are reported as best-fit values ± SE. Batch-to-batch reproducibility was assessed separately by comparing sensors from *N* = 3 independent fabrication runs performed on different days with separately prepared reagent solutions. The CV was calculated as SD/mean × 100; within-PPP CV was obtained from three reaction puddles on a single 96-PPP, and between-batch CV from the means of the three 96-PPPs. Comparisons with a control group were performed by one-way ANOVA with Dunnett’s post hoc test, and comparisons between two groups by unpaired t-test. Differences with p < 0.05 were considered statistically significant. All statistical analyses were performed using GraphPad Prism 11.0.0.

## Acknowledgments

This research was funded by the National Research Foundation of Korea (NRF) under Grant No. RS-2025-16068427. This work was also supported by the Ministry of Health and Welfare (KH140292) and the MSIT (Ministry of Science and ICT), Korea, under the ITRC (Information Technology Research Center) (IITP-2026-RS-2023-00258971).

