## Supporting Information for "Surfactant-Assisted Colorimetric Signal Enhancement in Paper-Based Glucose Sensing"

### Table of Contents


**Figure S1.** Optimization of the coating sequence for the colorimetric glucose assay ························3

**Figure S2.** Effect of buffer composition on the colorimetric glucose assay·································4

**Figure S3.** Dry-state absorbance analysis of the colorimetric glucose assay ······························5

**Figure S4.** SEM analysis of the paper substrate without and with deposited GOx/HRP/ABTS ············6

**Figure S5.** Hydrodynamic size distributions of GOx/HRP enzyme solutions without and with Tween 20·7

**Figure S6.** AFM analysis of GOx- and HRP-coated mica·······················································8

**Figure S7.** Effect of drying on glucose assay response and apparent reaction kinetics ······················9

**Figure S8.** Interference test of the glucose assay ··································································10

**Figure S9.** Glucose detection in artificial saliva ···································································11

**Figure S10.** Within-PPP and between-batch reproducibility of the glucose assay ···························12

**Figure S11.** TMB-based colorimetric glucose assay ·······························································13

**Table S1.** Four-parameter logistic (4PL) fitting parameters and analytical performance for each surfactant condition in Figure 2b ·······································································································14

**Table S2.** 4PL fitting parameters and analytical performance for each Tween 20 concentration in Figure 2d ·························································································································15

**Table S3.** 4PL fitting parameters and analytical performance for sensors prepared with 0.1 mg/mL GOx, without and with Tween 20, in Figure 3d····································································16

**Table S4.** Within-PPP and between-batch reproducibility of the colorimetric response without and with Tween 20. ············································································································17

**Table S5.** Analytical performance of the paper-based glucose sensor without and with Tween 20·········18

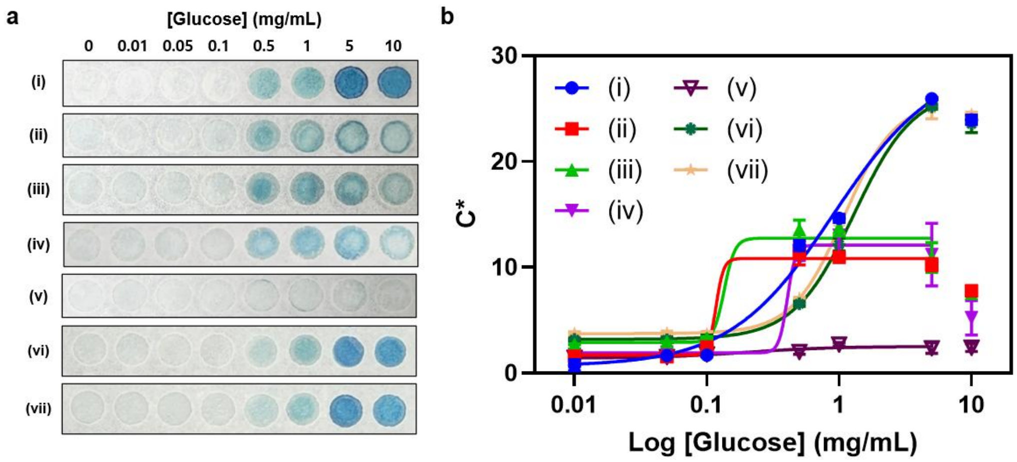


**Figure S1.** Optimization of the coating sequence for the colorimetric glucose assay. (a) Representative photographs and (b) calibration curves of the glucose assay prepared with different coating sequences of GOx, HRP, ABTS, and Tween 20. Conditions: (i) mixture (0.1 mg/mL GOx, 5 μg/mL HRP, 5 mM ABTS, 0.02% Tween 20); (ii) Tween 20 → ABTS → HRP → GOx (reverse order of molecular weight, excluding Tween 20); (iii) Tween 20 → GOx → HRP → ABTS (ascending molecular weight, excluding Tween 20); (iv) ABTS → Tween 20 → HRP → GOx (reverse molecular-weight order, including Tween 20); (v) GOx → HRP → Tween 20 → ABTS (ascending molecular-weight order, including Tween 20); (vi) Tween 20 → mixture (excluding Tween 20); (vii) ABTS → mixture (excluding ABTS). Data are presented as mean ± SD (*n* = 3).

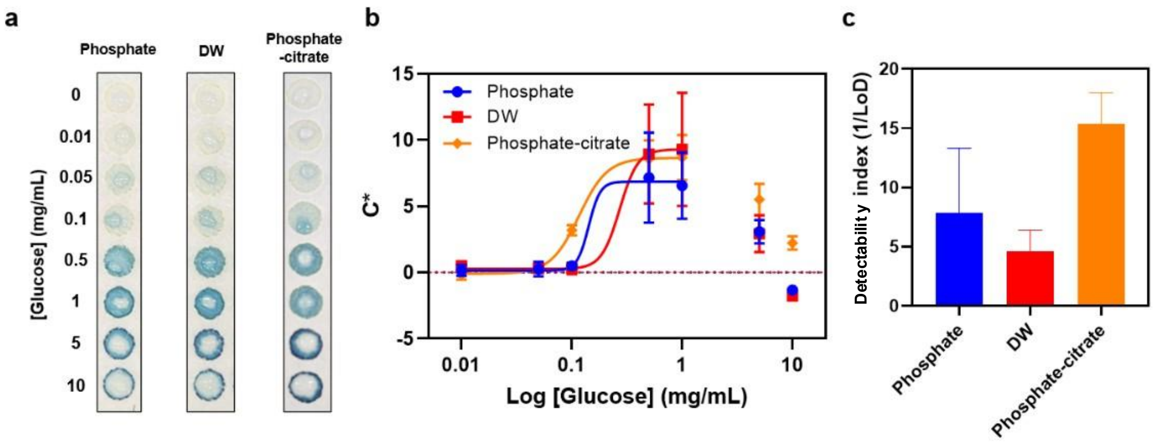


**Figure S2.** Effect of buffer composition on the colorimetric glucose assay. (a) Representative photographs, (b) calibration curves, and (c) detectability index (1/LoD ± SE) of the glucose assay prepared in phosphate-citrate buffer, phosphate buffer, or distilled water (DW). Sample volume: 10 µL per puddle. Data are presented as mean ± SD ($n$ = 3).

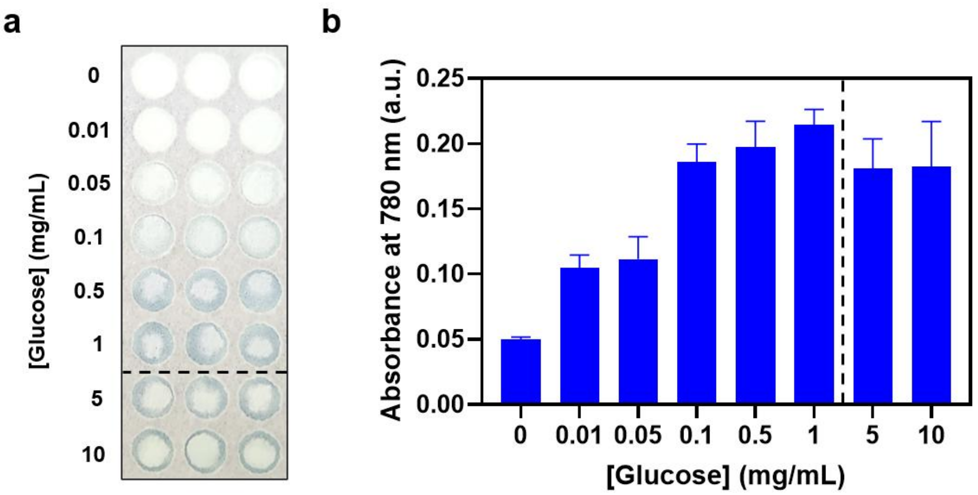


**Figure S3.** Dry-state absorbance analysis of the colorimetric glucose assay. (a) Photographs of GOx/HRP/ABTS-coated paper sensors after application of glucose solutions and subsequent drying, and (b) absorbance at 780 nm. Dashed lines separate the region where absorbance no longer increased with glucose concentration. Sample volume: 10 μL per puddle. Data are presented as mean ± SD ($n$ = 3).

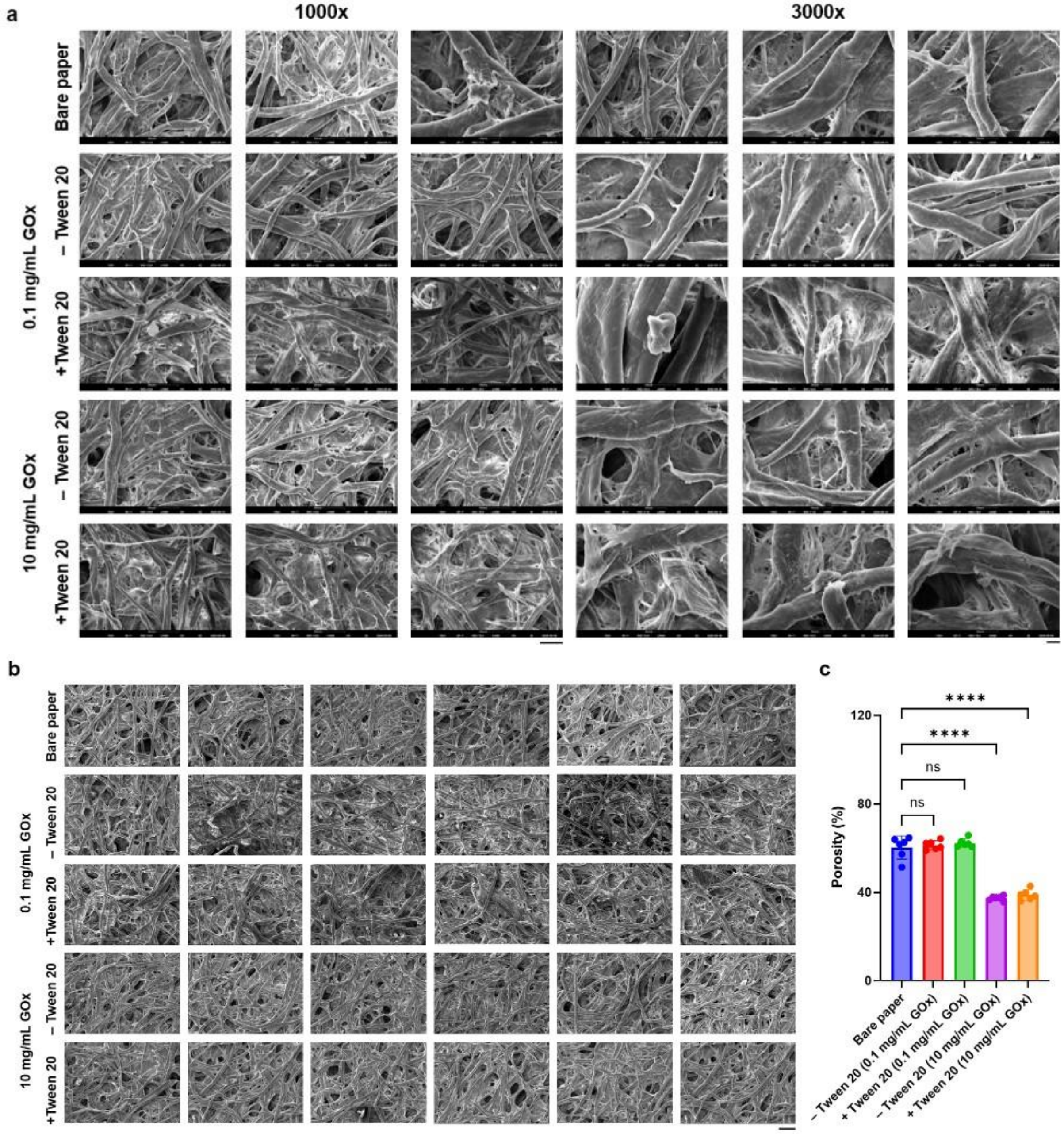


**Figure S4.** SEM analysis of the paper substrate without and with deposited GOx/HRP/ABTS. (a) SEM images of bare paper and sensors prepared at 0.1 and 10 mg/mL GOx, without and with Tween 20, acquired at 1000× (left three columns) and 3000× (right three columns). (b) Lower-magnification images (500×) of the same conditions used for porosity quantification. (c) Porosity determined from the images in (b). Data are presented as mean ± SD (*n* = 6); ns, not significant; ****$p < 0.0001$. Scale bars: 50 µm (a, left), 10 µm (a, right), 100 µm (b).

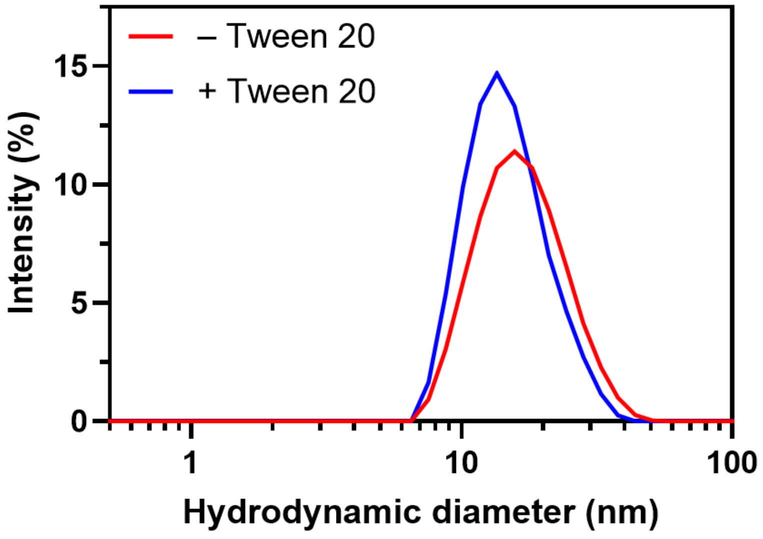


**Figure S5.** Hydrodynamic size distributions of GOx/HRP enzyme solutions without and with Tween 20. Distributions were measured by dynamic light scattering, and each represents the average of three repeated measurements of the same sample.

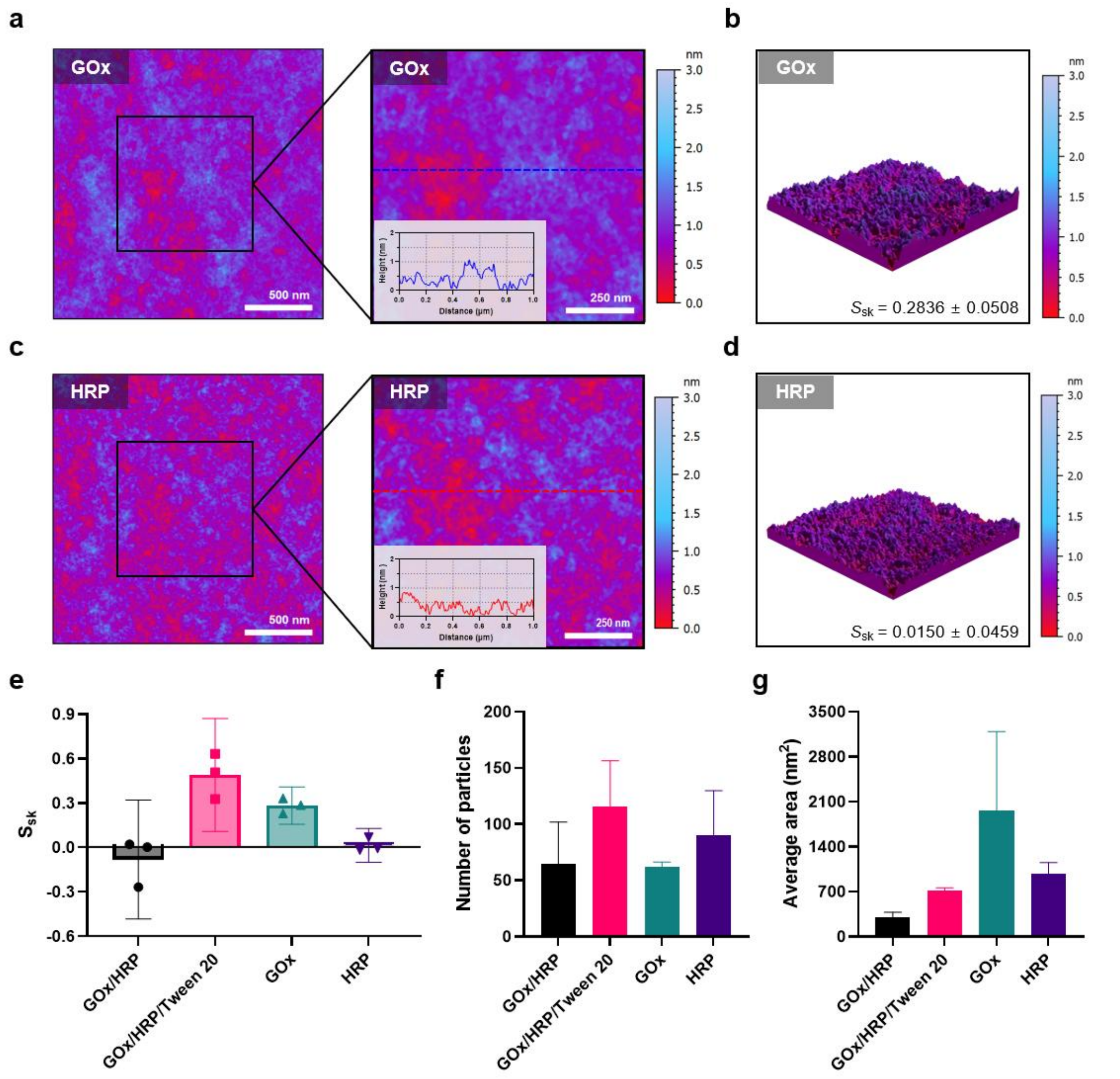


**Figure S6.** AFM analysis of GOx- and HRP-coated mica. (a) Height image of GOx-coated mica (2 × 2 μm$^2$, left) with a magnified image (1 × 1 μm$^2$, right); the inset shows the cross-sectional height profile along the dashed line in the magnified image. (b) 3D topographic image of the same surface. (c, d) Corresponding images for HRP-coated mica. (e) $S_{sk}$ values, (f) number of detected particles, and (g) particle areas for GOx/HRP, GOx/HRP/Tween 20, GOx, and HRP coatings. $S_{sk}$: surface skewness. All images are displayed on an identical z-scale (0–3 nm). Scale bars: 500 nm (full images), 250 nm (magnified images). Data are presented as mean ± SD ($n$ = 3).

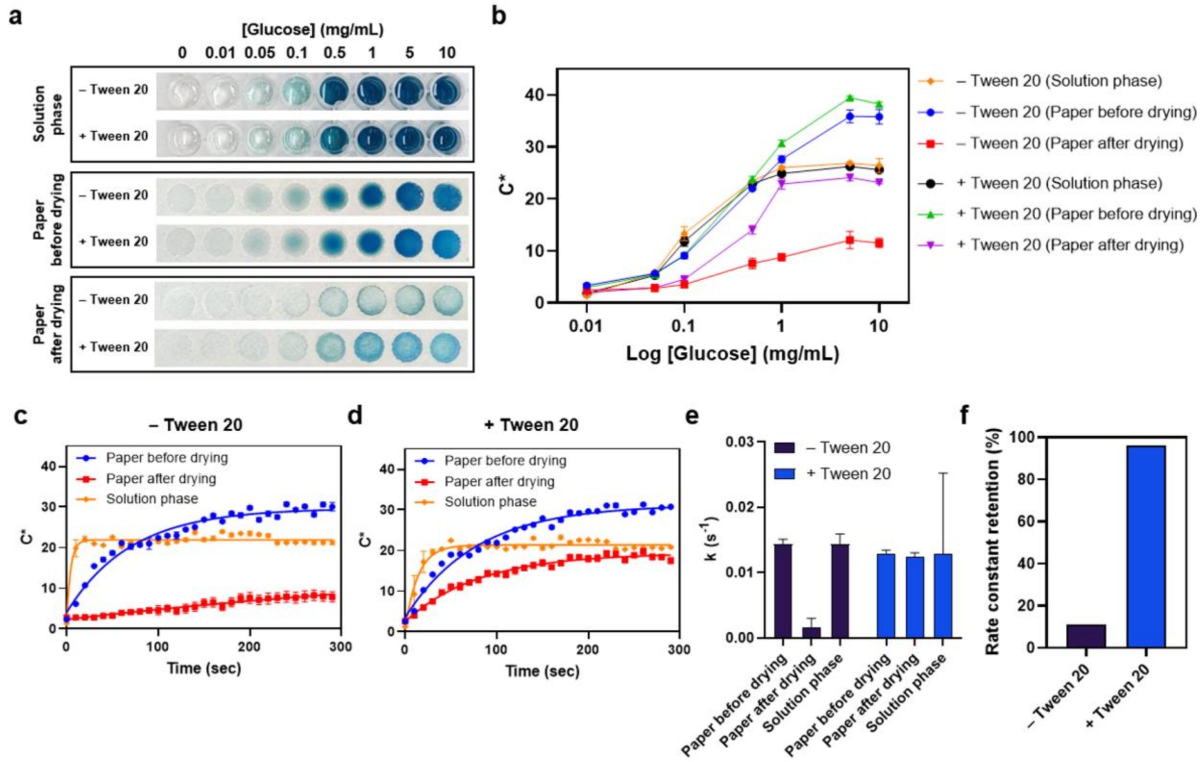


**Figure S7. Effect of drying on glucose assay response and apparent reaction kinetics.** (a) Representative colorimetric responses at different glucose concentrations in solution and on paper before and after drying. (b) $C^*$ as a function of glucose concentration under the indicated conditions. (c, d) Time-dependent $C^*$ responses without and with Tween 20. Data are presented as mean ± SD ($n$ = 3). (e) Apparent rate constants ($k$, $s^{-1}$) under the indicated conditions. Data are presented as best-fit value ± SE ($n$ = 3). (f) Rate-constant retention after drying without and with Tween 20.

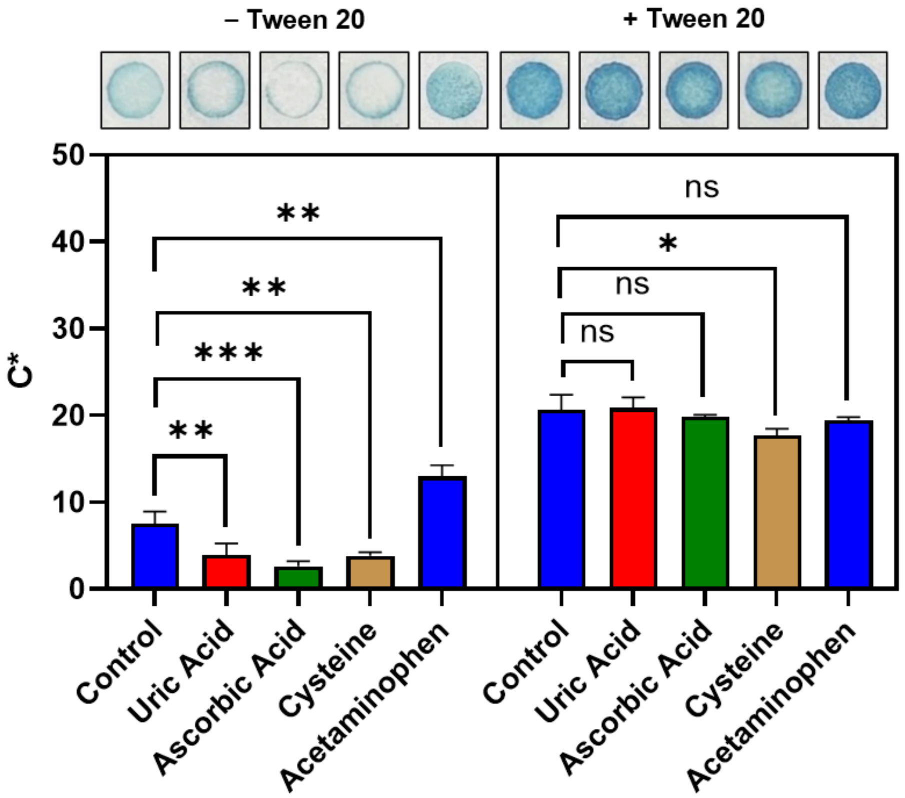


**Figure S8.** Interference test of the glucose assay. *C** responses to glucose in the presence of the indicated interfering compounds, without and with Tween 20. ns, not significant; *p < 0.05, **p < 0.01, ***p < 0.001, ****p < 0.0001. Data are presented as mean ± SD (*n* = 3).

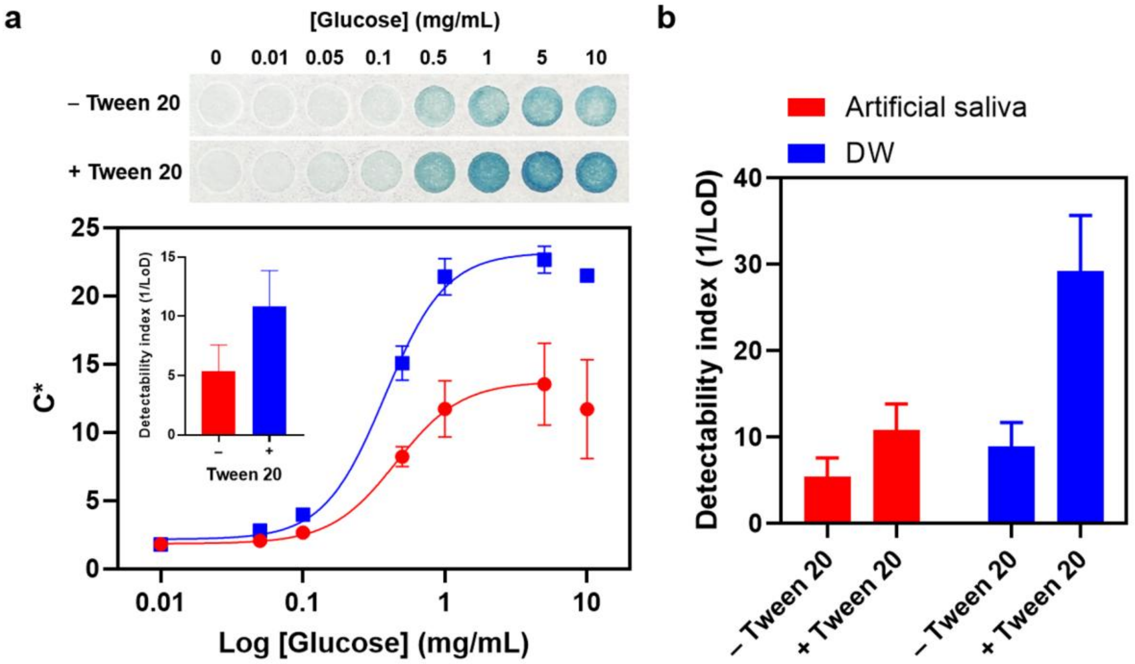


**Figure S9.** Glucose detection in artificial saliva. (a) Representative colorimetric responses and $C^*$ calibration curves in artificial saliva without and with Tween 20; inset, detectability index (1/LoD). Data are presented as mean ± SD ($n$ = 3). (b) Detectability index in artificial saliva and DW, without and with Tween 20. Data are presented as best-fit value ± SE ($n$ = 3).

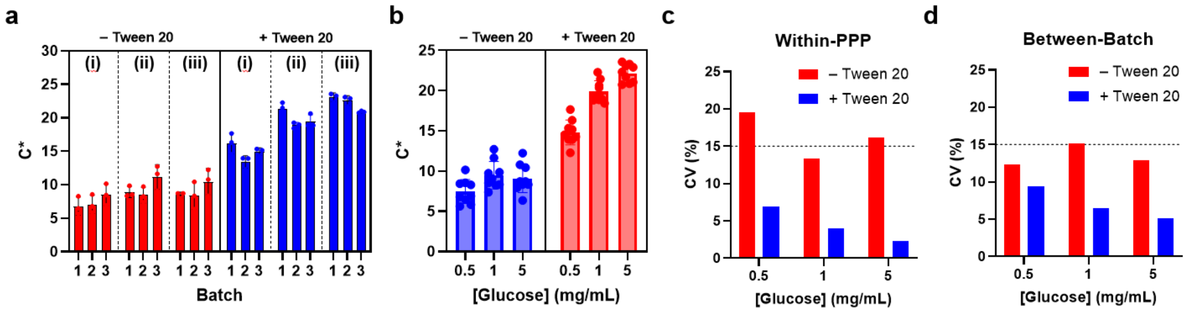


**Figure S10.** Within-PPP and between-batch reproducibility of the glucose assay. (a) $C^*$ responses of individual reaction puddles from three independently prepared 96-PPPs (batches 1–3) at (i) 0.5, (ii) 1, and (iii) 5 mg/mL glucose (mean ± SD, $n = 3$ puddles per batch). (b) $C^*$ responses pooled across the three batches (mean ± SD, $n = 9$). (c, d) Within-PPP and between-batch coefficients of variation (CV) at each glucose concentration. Dashed lines indicate the 15% CV benchmark commonly used for bioanalytical precision.[1]

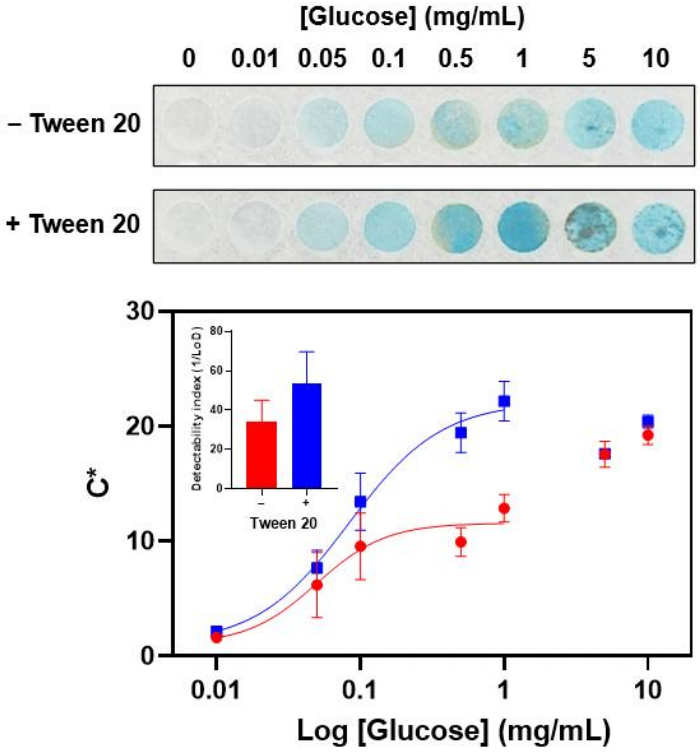


**Figure S11.** TMB-based colorimetric glucose assay. Representative colorimetric responses and corresponding *C** calibration curves in the absence and presence of Tween 20; inset, detectability index (1/LoD). Data are presented as mean ± SD (*n* = 3).

**Table S1**. Four-parameter logistic (4PL) fitting parameters and analytical performance for each surfactant condition in Figure 2b. Span and Hill slope were obtained from the 4PL fit, and $R^2$ indicates goodness of fit; the limit of detection (LoD) was interpolated from the fit as the glucose concentration corresponding to the bottom plateau plus three times its standard error.

| **Condition** | **Span** | **Hill slope** | $R^2$ | **LoD ± SE** |
|---|---|---|---|---|
| Control | 7.115 | 1.629 | 0.958 | 0.069 ± 0.018 |
| Tween 20 | 21.950 | 1.378 | 0.992 | 0.034 ± 0.007 |
| Tween 80 | 22.510 | 1.662 | 0.993 | 0.085 ± 0.020 |
| Triton X-100 | 31.140 | 1.193 | 0.988 | 0.068 ± 0.020 |
| SDS | 26.730 | 1.328 | 0.987 | 0.074 ± 0.020 |
| Dish detergent | 25.030 | 1.518 | 0.999 | 0.039 ± 0.008 |

**Table S2.** 4PL fitting parameters and analytical performance for each Tween 20 concentration in Figure 2d.

| [Tween 20] | Span | Hill slope | $R^2$ | LoD ± SE |
|---|---|---|---|---|
| 0% | 8.941 | 1.633 | 0.916 | 0.086 ± 0.022 |
| 0.02% | 19.830 | 1.441 | 0.995 | 0.034 ± 0.008 |
| 0.04% | 19.990 | 1.363 | 0.995 | 0.026 ± 0.007 |
| 0.06% | 17.870 | 1.383 | 0.993 | 0.038 ± 0.007 |

**Table S3.** 4PL fitting parameters and analytical performance for sensors prepared with 0.1 mg/mL GOx, without and with Tween 20, in Figure 3d.

| Condition | Span | Hill slope | $R^2$ | LoD ± SE |
|---|---|---|---|---|
| –Tween 20 | 7.347 | 2.132 | 0.933 | 0.113 ± 0.036 |
| + Tween 20 | 22.030 | 1.320 | 0.995 | 0.034 ± 0.008 |

**Table S4.** Within-PPP and between-batch reproducibility of the colorimetric response without and with Tween 20. Chroma values (*C**) were measured at three glucose concentrations; within-PPP variability was assessed from $n = 3$ reaction puddles on a single 96-PPP, and between-batch variability from $N = 3$ independently prepared plates fabricated on different days. Between-batch values were calculated from 96-PPP means and are therefore not directly comparable in magnitude to within-PPP values.

| [Glucose] (mg/mL) | –Tween 20 | | | | | + Tween 20 | | | | |
|---|---|---|---|---|---|---|---|---|---|---|
| | Mean (*C**) | Within-PPP | | Between-Batch | | Mean (*C**) | Within-PPP | | Between-Batch | |
| | | SD | CV (%) | SD | CV (%) | | SD | CV (%) | SD | CV (%) |
| 0.5 | 7.457 | 1.492 | 19.5 | 0.917 | 12.3 | 14.831 | 1.540 | 6.9 | 1.390 | 9.4 |
| 1 | 9.509 | 1.720 | 13.3 | 1.144 | 15.2 | 19.901 | 1.341 | 4.0 | 1.296 | 6.5 |
| 5 | 9.103 | 1.761 | 16.2 | 1.176 | 12.9 | 22.166 | 1.110 | 2.3 | 1.141 | 5.1 |

**Table S5.** Analytical performance of the paper-based glucose sensor without and with Tween 20.

| Parameter | − Tween 20 | + Tween 20 |
|---|---|---|
| **Detection performance** | | |
| LoD (mg/mL) | 0.113 ± 0.036 | 0.034 ± 0.008 |
| Detectability index (1/LoD) | 8.877 ± 2.813 | 29.216 ± 6.457 |
| Working range (mg/mL) | 0–5 | 0–5 |
| Span | 7.347 | 22.030 |
| **Reproducibility (5 mg/mL glucose)** | | |
| Within-PPP CV (%) | 16.2 | 2.3 |
| Between-batch CV (%) | 12.9 | 5.1 |
| **Functional retention after drying** | | |
| Rate constant retention after drying (%) | 11 | 95 |
| **Stability under stress** | | |
| Wetting–drying (k, $cycle^{-1}$) | 1.76 | 1.01 |
| Thermal stress (k, $min^{-1}$) | 0.09 | 0.04 |
| High ionic strength (% retained) | 26 | 77 |
| **Matrix performance** | | |
| LoD in artificial saliva (mg/mL) | 0.186 ± 0.076 | 0.092 ± 0.025 |
